# The altered entry pathway and antigenic distance of the SARS-CoV-2 Omicron variant map to separate domains of spike protein

**DOI:** 10.1101/2021.12.31.474653

**Authors:** Thomas P. Peacock, Jonathan C. Brown, Jie Zhou, Nazia Thakur, Ksenia Sukhova, Joseph Newman, Ruthiran Kugathasan, Ada W.C. Yan, Wilhelm Furnon, Giuditta De Lorenzo, Vanessa M. Cowton, Dorothee Reuss, Maya Moshe, Jessica L. Quantrill, Olivia K. Platt, Myrsini Kaforou, Arvind H. Patel, Massimo Palmarini, Dalan Bailey, Wendy S. Barclay

## Abstract

The SARS-CoV-2 Omicron/BA.1 lineage emerged in late 2021 and rapidly displaced the Delta variant before being overtaken itself globally by, the Omicron/BA.2 lineage in early 2022. Here, we describe how Omicron BA.1 and BA.2 show a lower severity phenotype in a hamster model of pathogenicity which maps specifically to the spike gene. We further show that Omicron is attenuated in a lung cell line but replicates more rapidly, albeit to lower peak titres, in human primary nasal cells. This replication phenotype also maps to the spike gene. Omicron spike (including the emerging Omicron lineage BA.4) shows attenuated fusogenicity and a preference for cell entry via the endosomal route. We map the altered Omicron spike entry route and partially map the lower fusogenicity to the S2 domain, particularly the substitution N969K. Finally, we show that pseudovirus with Omicron spike, engineered in the S2 domain to confer a more Delta-like cell entry route retains the antigenic properties of Omicron. This shows a distinct separation between the genetic determinants of these two key Omicron phenotypes, raising the concerning possibility that future variants with large antigenic distance from currently circulating and vaccine strains will not necessarily display the lower intrinsic severity seen during Omicron infection.

## Summary

The second year of the SARS-CoV-2 pandemic was marked by the repeated emergence of SARS-CoV-2 variants with altered antigenicity, and/or enhanced transmissibility. The fifth Variant Of Concern (VOC), Omicron, was first detected in Southern Africa^1^. Within just a month of its initial detection, Omicron had spread to account for the majority of SARS-CoV-2 cases globally^2^. The major Omicron lineages, BA.1 and BA.2 (as well as the emerging Omicron lineages BA.4 and BA.5), are marked by a total of >50 mutations across the entire genome, including an unprecedented >25 mutations in the spike glycoprotein gene^1,3,4^. The large antigenic distance between Omicron and the early pandemic spike protein correlates with a loss of neutralization titre, a lower vaccine effectiveness against Omicron, and a high frequency of reinfections^2,3,5-14^.

Infections with Omicron lineage viruses have resulted in a reduction in disease severity relative to the previous VOC, Delta, including in those previously uninfected, or unvaccinated individuals^15,16^. Omicron lineage isolates also show a lower intrinsic severity than previous VOCs in rodent models such as hamsters^17-20^. In *ex vivo* systems, Omicron shows attenuated replication in human cells derived from the lower respiratory tract but rapid replication in upper airway epithelia^21,22^. In vitro, Omicron does not induce syncytia, and the spike protein has low fusogenicity^12,19,23^. Syncytia have been reported at autopsy of COVID cases, and the efficient cleavage at the S1/S2 FCS in spike protein that underlies syncytia formation has been associated with enhanced disease severity in animal models^28,30,31^. Furthermore, Omicron has the ability to enter cells efficiently through the endosomal pathway compared to previous variants^12,19,21,23^. It is likely that these phenotypic properties are linked, and that cell entry route and fusogenicity are indicators/surrogates for severity.

Here we map the molecular basis of lower severity *in vivo* to the Omicron spike gene. We find that the Omicron spike confers attenuated replication in Calu-3 lung cells, due to endosomal restriction. However Omicron spike confers rapid replication in primary human nasal epithelial cultures, where endosomal restriction is less potent. Using chimeric and mutant recombinant spike proteins, we demonstrate that a single substitution in the S2 domain of Omicron spike, N969K, confers endosomal entry and leads to lower fusogenicity. Importantly, this means that the determinants of the cell entry pathway of Omicron (S2 domain) are distinct from those that confer its extreme antigenic distance (S1 domain). Thus, we emphasize that future variants with a similar antigenic distance to Omicron but no reduction in intrinsic severity could arise.

## Results

### The Omicron variants BA.1 and BA.2 show lower pathogenicity in a hamster model

To compare the intrinsic virulence of Omicron isolates with previous variants *in vivo*, we inoculated groups of at least 4 naïve hamsters intranasally with SARS-CoV-2 variants - WT/D614G (Pango lineage B.1), Alpha (B.1.1.7), Delta (B.1.617.2), Omicron/BA.1 (BA.1) or Omicron/BA.2 (BA.2) - and observed weight loss, a correlate of pathogenicity (Figure 1A). All animals were productively infected and shed infectious virus from the nose (Figure 1A, Extended data Figure 1A). Whereas those hamsters infected with WT(D614G), Alpha and Delta viruses showed a maximum mean weight loss between 9% and 14% in the first week post-infection. In contrast animals infected with either BA.1 or BA.2 gained weight during the experiment at a rate comparable to mock-infected animals. This confirms, as others have shown^17-20^, that BA.1 and BA.2 virus show an attenuated pathogenicity phenotype in the hamster model. We confirmed that pseudovirus (PV) expressing BA.1 spike efficiently enters cells expressing hamster ACE2, which shows that the lack of pathogenicity is not due to inefficient cell entry (Extended data figure 1B).

**Figure 1.**
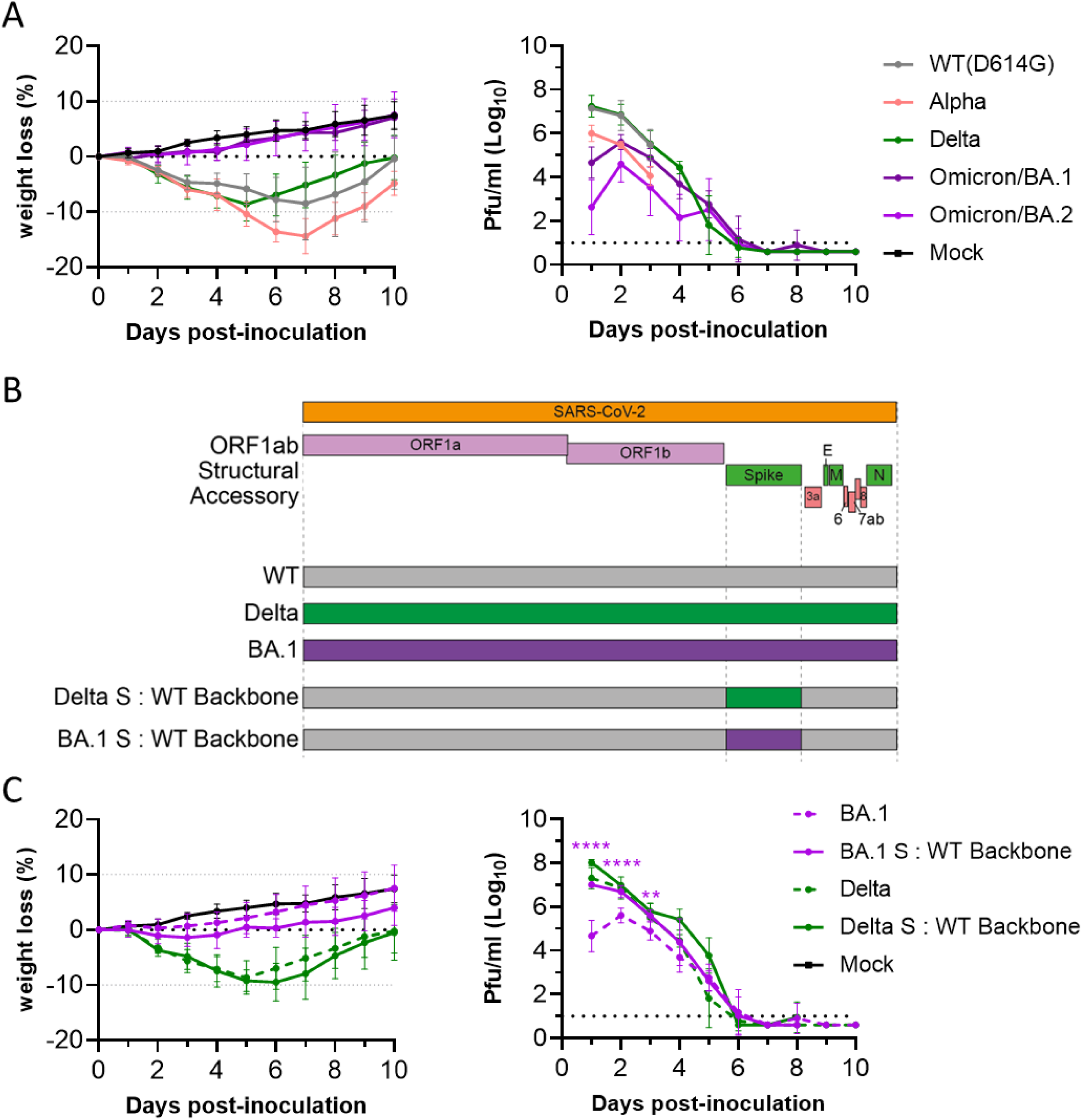
The pathogenicity phenotypes of SARS-CoV-2 variants Delta and Omicron map to the spike gene. (A,C) Weight loss and infectious virus shedding of hamsters directly inoculated intranasally with 100 plaque forming units (pfu) of the named variant or with PBS (Mock). Infectious virus shedding measured by plaque assays of nasal washes. At least 4 hamsters were included in each group. Hamsters were weighed and nasal-washed daily for 10 days. Dotted lines in part (C) indicate the same groups of hamsters included in part (A). (B) Schematic of reverse genetics spike chimeras used in this study containing the backbone from the reference strain Wuhan-Hu-1 (WT) and the spike gene from the named variant. Statistics in (C) were performed by multiple two-tailed t tests (on log transformed data for the right panel). **, P<0.01; ****, P<0.0001. Purple stars indicate significant differences between BA.1 and BA.1 S: WT Backbone.

### The pathogenicity phenotypes of both the Delta and Omicron BA.1 variants map to the spike protein

To map the genetic basis of this pathogenicity phenotype, we used recombinant viruses with the spike gene of Delta or Omicron/BA.1 in the genetic backbone of the reference strain, Wuhan-Hu-1 (Figure 1B) as described in our recent study^12^. Groups of 4 hamsters were inoculated intranasally with an equal dose of each virus. Weight loss and viral shedding were compared to that induced by the corresponding whole virus isolates used previously (Figure 1A). Hamsters infected with the recombinant viruses harbouring Delta or Omicron spike showed similar weight loss and shedding to the full isolates. This suggests that the pathogenicity phenotype maps largely to the spike gene in the hamster model (Figure 1C, Extended data Figure 1C). Animals infected with the BA.1 spike chimera lost marginally, though not significantly, more weight and shed significantly higher titres of infectious virus than the full BA.1 infected animals on days 1-3 post-inoculation, suggesting that viral genes other than spike may contribute to the attenuated pathogenicity phenotype of Omicron (Figure 1C, Extended data Figure 1C).

### Omicron is attenuated for replication in Calu-3 lung cells but replicates more rapidly than Delta in primary cultures of human nasal epithelial cells

To compare replication of Delta and Omicron (BA.1) *in vitro*, each variant was used to inoculate Calu-3 cells - an immortalised human lung cell line expressesing the SARS-CoV-2 entry factors ACE2 and TMPRSS2^24^ - or transwells of primary human nasal epithelial cells grown at air-liquid interface (hNECs). Vero E6 cells overexpressing human ACE2 and TMPRSS2 (Vero-AT) cells were also inoculated, and replication of the two variants in these cells was found to be comparable (Figure 2A). In Calu-3 cells the viral yields of Omicron were lower than for Delta across all time points, reflecting the pattern seen in hamsters. Interestingly, this was not the case for hNECs, whereby BA.1 showed a large early replication advantage, releasing apical virus with infectious titres approximately 100-fold higher than Delta by 24 hours post-infection (Figure 2A, Extended data Figure 2A). However, by 48- and 72-hours post-infection, viral yields were lower compared to Delta.

**Figure 2.**
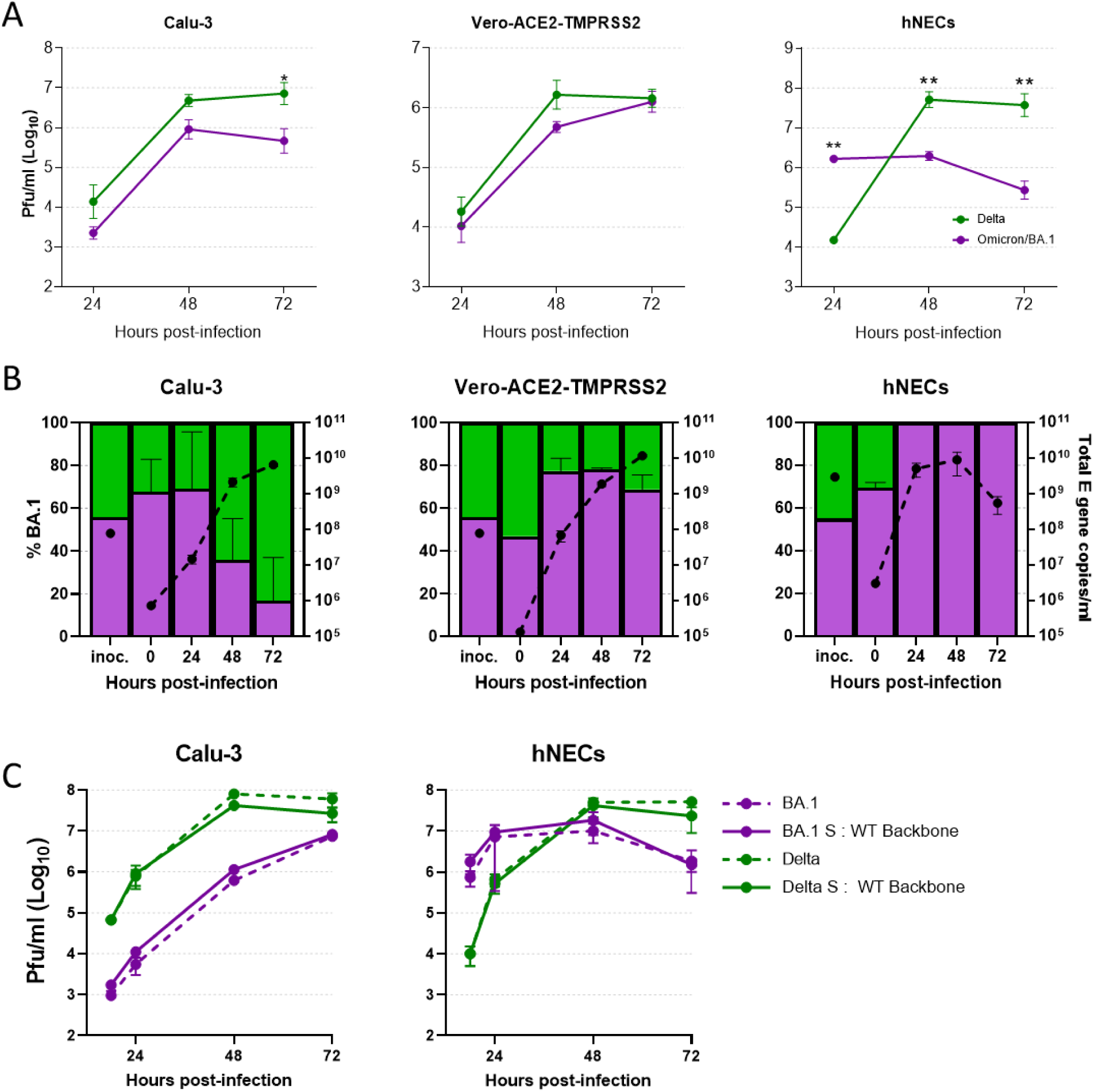
Comparative replication kinetics of Omicron and Delta. (A) Comparative replication kinetics of SARS-CoV-2 Omicron and Delta variants *in vitro*. Calu-3 cells or Vero-ACE2-TMPRSS2 (Vero-AT) were inoculated at an MOI of 0.001. Primary human nasal epithelial cultures (hNECs) were inoculated with virus at an MOI of 0.1. Infections were performed in triplicate and back titration confirmed equal inputs of the variants by RT-qPCR and plaque assay to measure E gene copies and infectious units, respectively. Assays were performed on at least 3 independent donors during 3 different weeks for primary cell experiments. Bars show individual replicate titres with the standard deviation. Statistical differences measured by ANOVA on log transformed data. *, P<0.05; **, P<0.01; ***, P<0.001; ****, P<0.0001. **–** (B) Competition assay between SARS-CoV-2 Omicron/BA.1 (purple) and Delta/B.1.617.2 (green) isolates *in vitro* and *ex vivo*. A probe-based RT-qPCR assay was developed to measure the proportion of Omicron and Delta present in mixed samples (Extended data Figure 3). The proportion of variant-specific signal is shown with the s.d. of 3 replicates. Equal mixtures of virus were inoculated onto hNECs at a final MOI of 0.1 and onto Vero-AT and Calu-3 cells at an MOI of 0.001. Daily harvests were measured for total viral load by E gene qRT-PCR (black dots) and by the variant specific RT-qPCR to give variant proportions. Mean + s.d. of 3 replicates shown. (C) Comparative replication kinetics of SARS-CoV-2 Omicron and Delta isolates and chimeric spike viruses *in vitro* as measured by release of infectious virus.

We also performed mixed competition assays in Vero-AT cells, Calu-3 cells or hNECs. Genome to RNA ratios of the two variants were measured over time following the inoculation of a 50:50 mix using a variant-specific PCR protocol (see Extended data Figure 3A-C). Again, in Calu-3 cells, BA.1 was attenuated and outcompeted by Delta. Genome copies of both viruses remained comparable in the Vero-AT cells across all time points, whereas in hNECs the BA.1 isolate rapidly outcompeted Delta (Figure 2B). We repeated the competition experiment using an independent set of Delta and Omicron isolates and found the same pattern of opposite replication phenotypes in the different cell systems (Extended Data Figure 2B).

We further compared the replication of Omicron/BA.1 and BA.2 isolates in hNECs and found BA.2 displayed the same initially rapid replication peaking at 24 hours (Extended data Figure 2C). In a direct competition assay between BA.1 and BA.2 in hNECs, neither virus outgrew the other, although BA.2 genome copies trended higher over the first 48 hours post-infection (Extended data Figure 2D).

### The attenuation of Omicron/BA.1 in Calu-3 and rapid replication in primary human nasal cells maps to the spike protein

To assess the role of the spike gene in determining the cell type-specific differences in replication between Omicron and Delta, we measured replication kinetics of the recombinant viruses that varied only in spike and compared them to their parental whole virus isolates. Each chimeric virus recapitulated the pattern of its spike donor parent, suggesting the spike gene is the major determinant of the attenuated replication of BA.1 in Calu-3 cells and its more rapid, but truncated, replication in hNEC cultures (Figure 2C, Extended data Figure 2E).

### Omicron spike protein shows enhanced human ACE2 binding but attenuated fusogenicity

The spike protein of SARS-CoV-2 acts as both the virus attachment protein, by binding to the host ACE2 receptor, and viral fusion protein, by facilitating the transfer of the genome to target cells. In addition the spike protein is the major antigen of the SARS-CoV-2 and primary immunogen in all currently licensed vaccines. Omicron spike carries several amino acid substitutions predicted to enhance binding to the ACE2 receptor^25^ and a number of groups have shown an approximately 2-3 fold increase in ACE2 binding compared with WT D614G spike receptor binding domain (RBD) or whole D614G spike protein^7,23,26^. We also measured an increase in the interaction between BA.1 spike and human ACE2 compared with the WT D614G spike binding (Extended data Figure 4A). Increased binding to the human ACE2 receptor might explain the faster replication we observed in human airway cells. However, it is unclear why, in alternative cell models, Omicron shows attenuated replication and results in less severe disease.

To mediate cell entry, spike protein must be activated through cleavage at the S2’ site by cellular proteases in order to enable fusion between the viral and host cell membranes. Ourselves and others have previously shown that, due to the presence of a furin cleavage site (FCS) at the S1/S2 junction, SARS-CoV-2 virions emerge from producer cells already primed for S2’ cleavage, and gain a replication and transmission advantage by rapidly entering airway cells at the cell surface or early endosome in a TMPRSS2-dependent manner^24,27^. A by-product of this fusion at the cell surface is syncytium formation, which is hypothesised to be correlated to disease severity^28-31^.

Omicron has been shown to lack the fusogenicity of previous variants such as Delta^12,19,23^. Indeed, we observe that infection of Vero-AT cells with 2 independent isolates of Omicron/BA.1 does not result in syncytia formation, in contrast to infection with WT/D614G or Delta virus isolates (Figure 3A).

**Figure 3.**
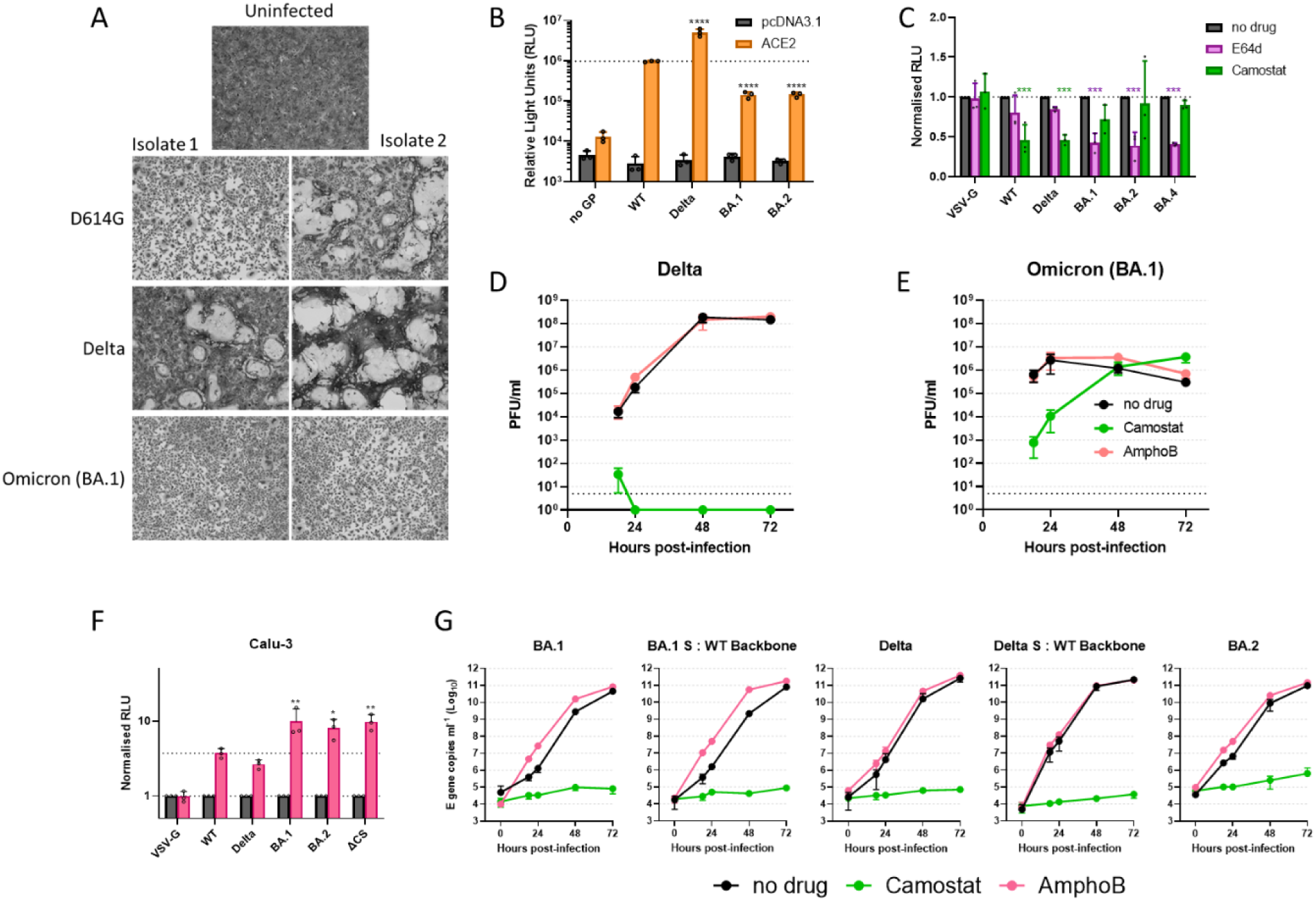
Cell entry mechanism of Omicron. (A) Live virus fusion assays of different SARS-CoV-2 variants. Vero-ACE2-TMPRSS2 cells were infected at an MOI of 0.25, left 18 hours, then fixed and stained with Giemsa solution. Results were repeated on two different occasions with representative fields shown. (B) Cell-cell fusion assays of BHK21 cells with rLUC-GFP1-7 expressing the stated spike protein and BHK-21 cells expressing human ACE2 ± TMPRSS2 and rLUC-GFP 8-11. All assays were performed in triplicate and are plotted as mean + s.d. One-way ANOVA with multiple comparisons performed on log-transformed data against WT. Representative graph from 3 independent repeats shown. (C, G) Inhibition/enhancement of pseudovirus expressing different viral glycoproteins into (C) Caco-2 or (F) Calu-3 cells. (C) The serine protease inhibitor, camostat (green), the cathepsin inhibitor, E64-d (dark pink) or (G) endosomal IFITM restriction factor inhibitor, Amphotericin B (bright pink) were used. Assays were performed in triplicate and are plotted as mean + s.d. Data shown is means values from three independent repeats (n = 3). Data was normalized to no drug control (grey). Statistical differences measured by Two-way ANOVA on (G only) log transformed data against (C) the no drug control, or (F) the WT plus amphotericin B control. *, P<0.05; **, P<0.01; ***, P<0.001; ****, P<0.0001. (D, E, G) Comparative replication kinetics of SARS-CoV-2 Omicron and Delta variants in (D,E) primary human nasal epithelial cultures (hNECs) or (G) Calu-3 inoculated with Delta or Omicron variants at an MOI (C) 0.1 or (D) 0.001. Cells were pre-treated both basolaterally and apically with either 50 µM of Camostat, 1µM of Amphotericin B or no drug for 2 hours prior to infection. 50 µM of Camostat or 1 µM Amphotericin B remained in the basolateral media for the course of the experiment. Infections were performed in triplicate wells and back titration confirmed equal inputs of the variants by E gene copies measured by RT-qPCR and infectious units measured by plaque assay. Plotted as mean ± s.d. of N=3 repeats. Experiments repeated on two separate occasions with representative repeat shown.

In a quantitative fusion assay, where spike proteins were expressed on the surface of cells and mixed with target cells expressing ACE2, neither BA.1, BA.2, nor BA.4 spike proteins mediated efficient cell fusion with target cells, whereas D614G or Delta spike drove extensive fusion (Figure 3B, Extended data Figure 4B,C). Thus, despite its increased ACE2 receptor binding strength, the cell surface fusion capacity of Omicron spike is attenuated compared to previous strains. This suggests Omicron spike is likely to mediate entry into cells through an alternative route.

### Omicron enters cells more efficiently through the endosomal route

In addition to cell surface/early endosomal entry via TMPRSS2, SARS-CoV-2 can also enter cells using the endosomal entry route wherein spike protein is cleaved and activated by endolysosomal proteases such as cathepsins. To investigate how SARS-CoV-2 spike variants mediate differential use of cell entry pathways, entry of PV bearing these spikes was measured in the presence of pathway-blocking drugs in Caco-2; a cell line we previously found permissive to both TMPRSS2 and cathepsin-mediated entry^24^. WT(D614G) and Delta spike-bearing PV were efficiently inhibited by Camostat (a broad inhibitor of serine proteases such as TMPRSS2) and poorly inhibited by E64d (a cathepsin inhibitor). This suggests that, in Caco-2, these PV enter at the cell surface or early endosome. In contrast, BA.1, BA.2 and BA.4 PV showed the opposite phenotype, being sensitised to E64d and refractory to Camostat inhibition (Figure 3C). This demonstrates that Omicron spike is better able to faciliate endosomal entry than previous variants, and that this property is conserved across all 3 major Omicron lineages (BA.1, BA.2 and BA.4/BA.5).

We also investigated the effect of Camostat on replication of authentic Omicron or Delta virus in hNEC (Figure 3D,E) or Calu-3 cells (Figure 3G). Camostat completely abrogated Delta virus replication in both cell lines, demonstrating a total dependence of Delta on serine proteases such as TMPRSS2. In contrast, the Omicron/BA.1 isolate replicated in hNECs in the presence of Camostat, albeit more slowly than in the absence of drug, eventually reaching equivalent titres by 48 hours post-infection (Figure 3D,E). This suggests that the higher early replication of Omicron in hNEC is TMPRSS2-mediated, although BA.1 can also efficiently use the endosomal pathway for replication (Extended data Figure 5A).

Single cell sequencing analysis of upper airway cells reveal there are approximately twice as many cells that are ACE2^+^/TMPRSS2^-^ than are ACE2^+^/TMPRSS2^+^ (Supplementary Table S1)^32-37^. We considered whether the early replication advantage of Omicron in the hNEC was conferred by an expansion of the number of target cells available for initiation of infection, as the endosomal pathway enables ACE2^+^/TMPRSS2^-^ cells to be infected. We modelled the comparative entry routes (Camostat sensitive vs Camostat insensitive) and established the enhanced that endosomal entry of Omicron cannot account for the more rapid replication in hNECs. Instead, Omicron appears to also use the TMPRSS2-dependent route more efficiently than Delta in this cell type (Supplementary Table S2, Extended data Figure 6A-D).

Ourselves and others have previously shown that entry through endosomes sensitizes SARS-CoV-2 viruses to inhibition by endosomal IFITM restriction factors, such as IFITM2/3^24,38^. However, this restriction can be lifted by the addition of Amphotericin B (AmphoB). Indeed, entry of BA.1 and BA.2 PV, as seen for PV with the ΔCS spike, a mutant spike without the furin cleavage site that is restricted to endosomal entry^24^, was significantly enhanced by AmphoB, whereas the entry of PV bearing other variants’ spike proteins was not (Figure 3F).

AmphoB treatment also enhanced yields of BA.1 or BA.2, but not Delta titres, in Calu-3 (Figure 3G). Using the recombinant chimeric spike-bearing viruses, we demonstrated this again mapping with mapping to the spike gene. Interestingly, in hNECs, AmphoB did not affect virus yields of Omicron (Figure 3D,E), suggesting that in upper airway cells Omicron is less inhibited by IFITM proteins. We previously showed considerable enhancement of virus replication of SARS-CoV-2 mutant lacking the FCS (ΔCS) that entered cells exclusively via the endosome following AmphoB treatment in airway cultures^24^. The substitutions in Omicron spike enable the virus to productively utilize the endosomal pathway in hNECs, whilst avoiding endosomal IFITM restriction.

### The altered entry route and antigenic distance of SARS-CoV-2 Omicron map to separate domains of the spike protein

To investigate the molecular basis of the altered entry route and low fusogenicity of Omicron variants, we engineered spike protein chimeras between WT(D614G) and Omicron/BA.2. McCallum *et al* recently identified several residues in the S2 domain that introduce additional electrostatic contacts within and between the spike protomers of BA.1 that seem likely to stabilize the protein and potentially impede fusion^39^. In this region four mutations are conserved between BA.1, BA.2 and BA.4^1,4^, which we found all shared a preference for the endosomal entry route in Caco-2 (Figure 3C). Therefore, we produced chimeric spike expression constructs with breakpoints after the S1/S2 cleavage site (containing the 4 common mutations N764K, D796Y, Q954H and N969K; Figure 4A). PV expressing chimeric spike protein, harbouring the S2 of BA.2, predominantly entered via the endosomal route, whereas the WT S2 domain conferred TMPRSS2-dependent cell surface entry to a spike with BA.2 S1 (Figure 4B). This pattern was also partially reflected in the syncytia formation assay, wherein lower fusogenicity was determined by the S2 domain of BA.2 (Figure 4C, D, Extended data Figure 5B), consistent with a recent report from Willett *et al*^12^. We further mapped the entry phenotype, and fusion phenotype to the single amino acid substitution, N969K (Figure 4B,D). PV bearing WT spike with the Omicron-like mutation N969K exhibited cathepsin-dependent entry similar to full BA.2 spike PV, whereas PV bearing BA.2 spike with the sole reversion K969N lost cathepsin sensitivity and instead became Camostat sensitive, similar to WT spike-bearing PV (Figure 4B). Interestingly, analysis of the extent of S1/S2 cleavage of spike in PV revealed that Omicron (BA.1, BA.2 and BA.4) spikes were efficiently cleaved with almost no full length spike detectable in concentrated PV (Figure 4E). This is contrary to several previous reports^19,23^, but in line with a recent report from Mesner *et al*^40^. This also correlates with single amino acids changes adjacent to the FCS in Omicron - specifically N679K and P681H - that on their own enhance cleavage^41^ (Extended data figure 5C,D). Substitutions in S2 including residue 969 did not impact spike FCS cleavage (Figure 4E).

**Figure 4.**
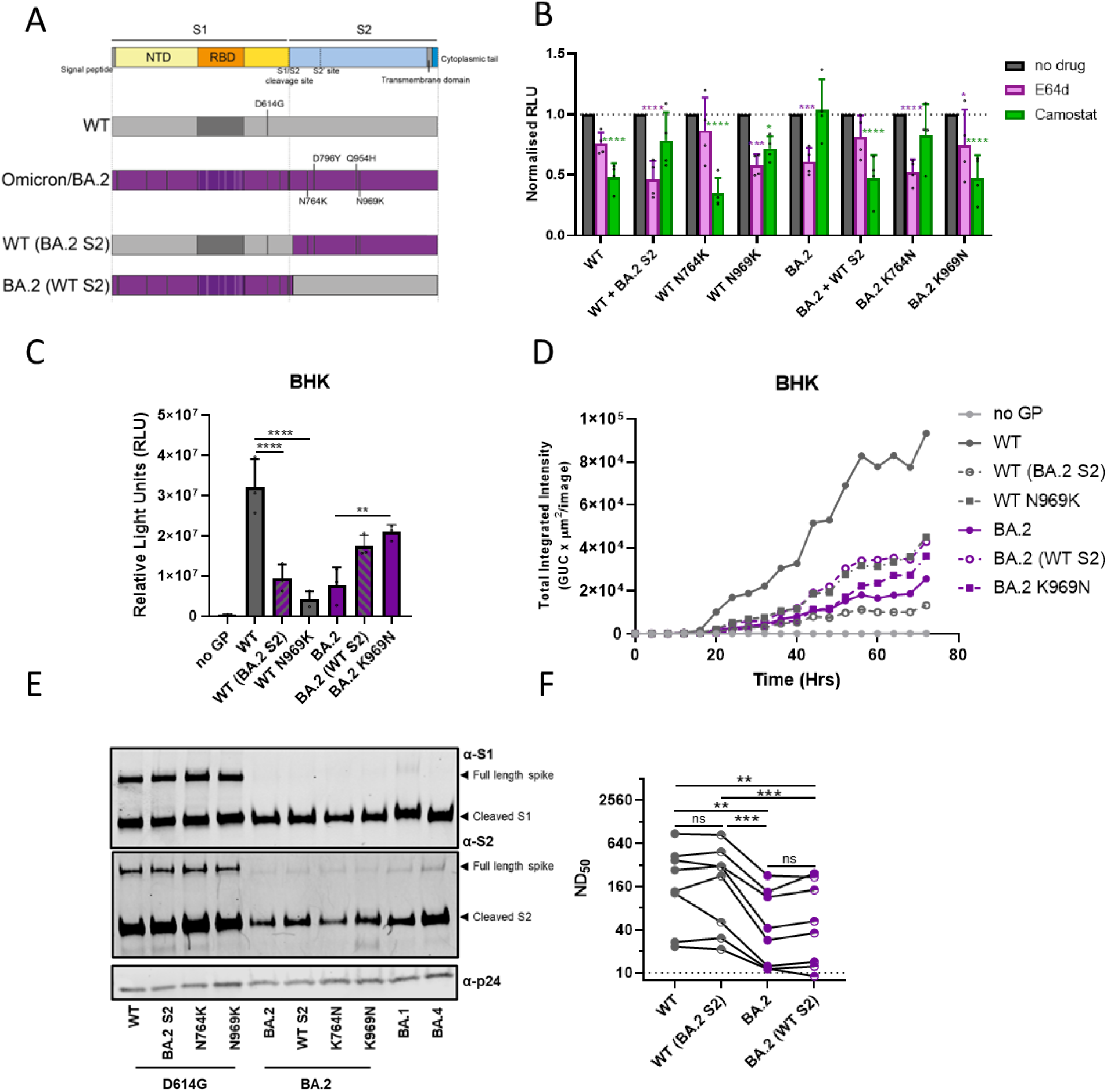
The enhanced efficiency of endosomal entry by Omicron maps to the spike S2 domain and is distinct from the determinant of antigenicity. (A) Schematic of spike Chimeras used to map out the enhanced endosomal entry phenotype of BA.2. (B) Inhibition of entry of pseudoviruses expressing different viral glycoproteins into Caco-2 the serine protease inhibitor, camostat (green), or the cathepsin inhibitor, E64-d (pink). Assays were performed in triplicate. Data shown are the means + s.d from four independent repeats (n = 4). All data were normalized to no drug control (grey). Statistical differences measured by Two-way ANOVA with multiple comparisons against the no drug control. *, P<0.05; **, P<0.01; ***, P<0.001; ****, P<0.0001. (C, D) Cell-cell fusion assays of BHK21 cells with rLUC-GFP1-7 expressing the stated spike protein and BHK-21 cells expressing human ACE2 and rLUC-GFP 8-11. All assays were performed in triplicate and are plotted as mean + s.d. Representative repeat of N=3 independent repeats shown. One-way ANOVA with multiple comparisons was performed. *, P<0.05; **, P<0.01; ***, P<0.001; ****, P<0.0001 (F) Neutralisation assay of the different spike chimera-containing pseudoviruses with 2 dose vaccine antisera (N=8). Representative repeat of N=3 independent repeats shown. Statistics performed on log2 transformed data with One-Way ANOVA with multiple comparisons. **, P<0.01; ***, P<0.001

The rapid spread of Omicron can be accounted for by a combination of a potential increase in inherent transmissibility, but also a large antigenic distance from previous variants and the spike protein of the vaccines that have now been widely used, allowing frequent re-infections and vaccine-breakthrough infections^42^. The antigenic determinants of spike protein have been mapped largely to the S1 domain, with epitopes both in the NTD and the RBD^43^. To demonstrate that the antigenicity of the S1/S2 chimeras was determined by the sequence in S1, we performed PV neutralisation assays with post-second dose vaccine antisera (Figure 4F). As expected, the S2 domain had no effect on the antigenic properties of the spike proteins, confirming that the entry/fusion phenotype and the antigenic phenotypes of Omicron map to discrete spike substitutions.

## Discussion

Omicron has shown one of the most rapid global takeovers of any SARS-CoV-2 variant to date. Its rapid growth derives from a combination of partial escape from prior immunity, and the phasing out of non-pharmaceutical interventions in many countries that coincided with its emergence. Omicron appears to be inherently more transmissible than previous variants^42^, with a shorter serial interval than Delta, but is also associated with less severe disease. A key issue is to establish whether these three traits of antigenic distance, high transmissibility and lower severity are linked, and this would have the implication of an inevitable attenuation of severity for future variants. Here, we attribute these phenotypes to distinct regions within the spike gene, suggesting they are likely to diverge as the virus continues to evolve beyond Omicron.

We show that the altered entry route of Omicron is partly conferred by substitutions in the S2 domain, particularly the substitution N969K that likely stabilises the interactions between protomers and hinders fusion^39^. This slows altered entry and reduces cell surface fusion, a process that has been linked with disease severity. Lack of syncytia formation by Omicron, despite high ACE2 receptor binding and efficient S1/S2 cleavage, may be accounted for by the slowed fusion conferred by S2 mutations. Moreover, in lung cells, where innate immune responses are potent^44^, slower entry through the late endosomes renders virus more vulnerable to restriction factors, and the resulting attenuation of replication also likely contributes to the less severe disease outcome. In contrast, the innate response of cells of the upper airway is dampened^44^ and the slower entry does not compromise replication. One early hypothesis for increased transmission of Omicron was that the numbers of susceptible cells are increased by the ability of Omicron to enter cells that do not express TMPRSS2. Here, we disprove this hypothesis by using modelling to show that the rapid replication in primary nasal cells is still largely TMPRSS2-dependent.

Although we have shown the lower severity phenotype of Omicron largely maps to the spike protein, consistent with a recent report by Barut *et al*^22^, further attenuating mutations may exist in Omicron outside this gene. For example, the recently emerged BA.1 (spike) x Delta recombinant XD, which contains most of the spike gene of BA.1 with the rest of its genome from Delta, is more virulent than whole Omicron^45^, and a recombinant virus bearing a BA.2 spike on a Wuhan-Hu-1 backbone showed higher pathogenicity in a hamster model^46^. Further mapping using reverse genetics will be required to identify the exact contribution of non-Spike mutations towards Omicron pathogenicity.

Antigenic distance is encoded for by substitutions in the major epitopes that lie within the S1 region of spike protein. This region also encodes enhanced ACE2 binding of Omicron and may be partially responsible for the higher transmissibility of Omicron. Conversely, the slower entry, and lower fusogenicity of Omicron largely maps to the S2 domain of spike. In recent months several Delta x BA.1 recombinants have been described, including some with breakpoints within spike^45,47^. Although not yet detected, we suggest a recombinant containing the S1 domain of Omicron (BA.1 or BA.2) and the S2 domain of Delta (or a different non-Omicron variant) may have the antigenic escape properties of Omicron, and entry route, and therefore, perhaps the intrinsic severity phenotype of Delta. Utmost efforts must be made to enable rapid identification of the emergence of such recombinants.

In conclusion, this work suggests that future variants could arise with distant antigenic properties but Delta-like severity phenotypes as both properties map to discrete regions of the spike protein. Whilst the circulation of SARS-CoV-2 remains elevated in many regions of the world, the possibility for future waves of severe disease cannot be dismissed.

## Materials and methods

### Biosafety and ethics statement

All laboratory work was approved by the local genetic manipulation safety committee of Imperial College London, St. Mary’s Campus (centre number GM77), and the Health and Safety Executive of the United Kingdom, under reference CBA1.77.20.1. SARS-CoV-2 reverse genetics work was performed at CVR University of Glasgow under HSE GM notification number is GM223/CVR_MP_AP amd3. Animal research was carried out under a United Kingdom Home Office License, P48DAD9B4.

Post-vaccine antisera samples were taken in concordance with the World Medical Association’s Declaration of Helsinki. The use of these sera was approved by the CDRTB Steering Committee in accordance with the responsibility delegated by the National Research Ethics Service (South Central Ethics Committee Oxford – C, NRES references 15/SC/0089 and 20/SC/0226).

### Hamster pathogenicity studies

Hamster pathogenicity studies were performed in a containment level 3 laboratory, using ISO Rat900 Individually Ventilated Cages (IVC) (Techniplast, U.K). Outbred Syrian Hamsters (4-7 weeks old), weighing 70-140 g were used. Prior to the study hamsters were confirmed to be seronegative against SARS-CoV-2. At least 4 hamsters per variant group were intranasally inoculated with 50 μl of 100 PFU/hamster of the named virus while lightly anaesthetised with isoflurane. All animals were nasal washed daily by instilling 400 μl of PBS into the nostrils and the expectorate collected into disposable 50 ml falcon tubes. Hamsters were weighed daily post-infection.

### Cells

Human embryonic kidney cells (293T; ATCC CRL-11268) were maintained in complete media (DMEM, 10% FCS, 1% non-essential amino acids (NEAA) and 1% penicillin-streptomycin (P/S)). African green monkey kidney cells overexpressing human ACE2 and TMPRSS2 (Vero E6-ACE2-TMPRSS2; Glasgow University)^48^, were maintained in complete media supplemented with 200 µg/ml hygromycin B (Gibco) and 2mg/ml G418 (Gibco) to maintain human ACE2 and TMPRSS2 expression. Human lung cancer cells (Calu-3; ATCC HTB-55) and Human epithelial colorectal adenocarcinoma cells (Caco-2; ATCC HTB-37) were maintained in DMEM, 20% FCS, 1% NEAA and 1% P/S. Baby hamster kidney cells (BHK-21; ATCC CCL-10) were maintained in DMEM supplemented with 10% FCS, 1 mM sodium pyruvate solution (Sigma-Aldrich, Germany), and 1× P/S. Air–liquid interface human nasal epithelial cells (hNECs) were purchased from Epithelix and maintained in Mucilair cell culture medium (Epithelix). All Cells were maintained at 5% CO_2_, 37°C.

### SARS-CoV-2 virus and reverse genetics

Upper respiratory tract swabs used to isolate viruses were collected for routine clinical diagnostic use and sequenced using the ARTIC network protocol (https://artic.network/ncov-2019) to confirm the presence of BA.1 lineage virus, under approval by the Public Health England Research Ethics and Governance Group for the COVID-19 Genomics UK consortium (R&D NR0195). Delta and Omicron virus were isolated by inoculating 100 µL of neat swab material onto Vero E6-ACE2-TMPRSS2 cells, incubating at 37°C for 1 h before replacing with growth media supplemented with 1 x penicillin/streptomycin and 1 x amphotericin B. Cells were incubated for 5-7 days until cytopathic effect was observed. Isolates were passaged a further two times in Vero E6-ACE2-TMPRSS2 cells^48^.

Virus growth kinetics and competition assays were performed as described previously^24^. Briefly, in air-liquid interface hNECs, before infection cells were washed with serum-free media to remove mucus and debris. Cells were infected with 200 µL of virus-containing serum-free DMEM and incubated at 37°C for 1 h. Inoculum was removed, and cells were washed twice with PBS. Time points were taken by adding 200 µL of serum-free DMEM and incubating for 10 mins and 37°C before removal and titration.

For SARS-CoV-2 plaque assays, serial dilutions of virus supernatant in serum-free DMEM, 1% NEAA and 1% P/S were performed and inoculated onto VAT cells for 1 h at 37 °C. Inoculum was then removed and replaced with SARS-CoV-2 overlay medium (1 × MEM, 0.2% w/v BSA, 0.16% w/v NaHCO3, 10 mM HEPES, 2 mM L-Glutamine, 1 × P/S, 1% w/v Avicel). Plates were incubated for 3 days at 37 °C before overlay was removed and cells were stained for 1 h at room temperature in 2 x crystal violet solution.

Reverse genetics derived viruses were generated as previously described^12,52,53^. Briefly, recombinant SARS-CoV-2 Wuhan cDNA genomes bearing the D614G, or Delta spike- or Omicron spike-encoding sequence were generated from a set of relevant overlapping genomic cDNA fragments and assembled using the Transformation-Associated Recombination (TAR) in yeast method as described^12^. The assembled genomes were then used as a template to *in vitro* transcribe viral genomic RNA which was then transfected into BHK-hACE2-N cells stably expressing the SARS-CoV-2 N and human ACE2 gene^48^ for virus rescue. The rescued viruses were passaged once (P1 stock) in VERO E6 cells and their full genomes sequenced using Oxford Nanopore as previously described^54^, to confirm the presence of the desired mutations and exclude the presence of other spurious mutations.

### Plasmids, pseudovirus, pseudovirus entry and neutralisation assays

Spike-encoding pcDNA3.1 plasmids were generated by mutagenesis or by gene synthesis as described elsewhere^5,49^. Pseudovirus was generated and concentrated as previously described^24^. All spike expression plasmids used in this study contain D614G and K1255*STOP (that results in deletion of the C-terminal cytoplasmic tail of spike containing the endoplasmic retention signal, aka the Δ19 spike truncation).

Species ACE2 entry assays were performed as previously described^55^. Briefly, BHK cells were transfected with 500 ng of ACE2 or empty vector (pDISPLAY) using TransIT-X2 (Mirus Bio) according to the manufacturer’s recommendation. 24 h later, media was removed, and cells were harvested following the addition of 2mM EDTA in PBS, resuspended in DMEM and plated into white-bottomed 96 wells plates (Corning). Cells were overlayed with pseudovirus and incubated for 48 h. Firefly luciferase was quantified whereby media was replaced with 50 µL Bright-Glo substrate (Promega) diluted 1:2 with PBS and read on a GloMax Multi+ Detection System (Promega).

Pseudovirus neutralisation assays were performed by incubating serial dilutions of heat-inactivated human post-vaccine antisera with a constant amount of pseudovirus. Antisera/pseudovirus mix was then incubated at 37°C for 1 h and then had HEK 293T-ACE2 cells overlayed onto them. 48 h later assays were then lysed and read on a FLUOstar Omega plate reader (BMF Labtech) using the Steady-Glo^®^ luciferase assay system (Promega)

### Cell-cell fusion assays

Cell-cell fusion assays were performed as described elsewhere^55,56^. Briefly HEK 293T rLuc-GFP 1–7 effector cells were transfected in OptiMEM (Gibco) using Transit-X2 transfection reagent (Mirus), with 500 ng the named SARS-CoV-2 spike or mock transfected with empty pcDNA3.1. BHK-21 stably expressing rLuc 8-11 co-transfected with 500 ng of human ACE2. After 24 h, cells were harvested using 2 mM EDTA in PBS and 5×10^4^ effector and target cells in 100 µL volume were co-cultured. Quantification of cell–cell fusion was measured based on Renilla luciferase activity, 24 h later by adding 1 μM of Coelenterazine-H (Promega) at 1:400 dilution in PBS. The plate was incubated in the dark for 2 min and then read on a Glomax Multi+ Detection System (Promega) as above. GFP fluorescence images were captured every 3 h for 72 h using an Incucyte S3 real-time imager (Essen Bioscience, Ann Arbor, Michigan, USA). Five fields of view were taken per well at 10x magnification, and GFP expression was determined using the total integrated intensity metric included in the IncuCyte S3 software (Essen BioScience). To analyse images generated on the IncyCyte S3, a collection of representative images is first taken to set fluorescence and cellular thresholds, which allows for the removal of background fluorescence and selection of cell boundaries (‘objects’) by creating ‘masks’. Following this, the total integrated intensity metric can be accurately calculated by the software, which takes the total sum of fluorescence, expressed as green count units (GCU) µm^−2^. Assays were set up with 3 biological replicates for each condition, and each experiment was performed 3 independent times.

### Whole virus cell-cell fusion assays

Vero-ACE2-TMPRSS2 cells were infected with different SARS-CoV-2 variants/mutants at an MOI of 0.25. After 2 h, the inoculum was replaced with DMEM, 10% FBS. At 18 h post-infection, cells were fixed with 4% PFA, stained with Giemsa (SLS) and images were taken using the Evos FL Auto 2 microscope (Invitrogen).

### Flow cytometry-based spike-ACE2 affinity assay

HEK 293T cells were transfected with mammalian expression plasmids encoding the relevant SARS-CoV2 spike protein. At 24 h post-transfection, cells were dissociated and incubated with recombinant human ACE2-Fc(IgG) (1 mg/ml) (Abcam ab273687) at a dilution of 1:32000 for 30 minutes. Cells were then incubated with Goat Anti-Human IgG Fc (DyLight^®^ 650) preadsorbed (ab98622) for 30 minutes before determination of median fluorescence on a BD LSR Fortessa cell analyser.

### Western Blotting

Virus or pseudovirus concentrates were lysed in 4x Laemmli buffer (Bio-rad) with 10% β-mercaptoethanol and run on SDS-PAGE gels. After semi-dry transfer onto nitrocellulose membrane, samples were probed with mouse anti-p24 (abcam; ab9071) and rabbit anti-SARS spike protein (NOVUS; NB100-56578) or rabbit anti-S1 spike protein (SinoBiological; 40591-T62). Near infra-red (NIR) secondary antibodies, IRDye^®^ 680RD Goat anti-mouse (abcam; ab216776) and IRDye^®^ 800CW Goat anti-rabbit (abcam; ab216773) were subsequently used to probe membranes. Western blots were visualised using an Odyssey DLx Imaging System (LI-COR Biosciences).

### Viral RNA extraction and E gene qPCR

Virus genomes were quantified by E gene RT-qPCR as previously described^57^. Viral RNA was extracted from cell culture supernatants or hamster nasal washes using the QIAsymphony DSP Virus/Pathogen Mini Kit on the QIAsymphony instrument (Qiagen). RT-qPCR was then performed using the AgPath RT-PCR (Life Technologies) kit on a QuantStudio™ 7 Flex Real-Time PCR System with the primers specific for SARS-CoV-2 E gene^58^. For absolute quantification of E gene RNA copies, a standard curve was generated using dilutions viral RNA of known copy number. E gene copies per ml of original virus supernatant were then calculated using this standard curve.

To measure the proportions of Omicron and Delta RNA from mixed samples a modified RT-qPCR protocol was developed using probes specific for the S gene of Omicron or Delta variants. Flanking primers that were able to recognise a region of both Delta and Omicron were used (Forward: TGGACCTTGAAGGAAAACAGGG and Reverse: TGGTTCTAAAGCCGAAAAACCC). A pair of probes were then designed - a FAM probe specific to a 9 base pair insertion/deletion in the Omicron S gene at amino acid position 214 (TTATAGTGCGTGAGCCAGAAGA), and a HEX probe specific to the same region of the Delta S gene sequence lacking this insertion/deletion (CTATTAATTTAGTGCGTGATCT). Alternatively, a second HEX probe was designed specific to this region of BA.2, which also lacks the BA.1-specific insertion/deletion but has a SNP in this region (CTATTAATTTAGGGCGTGATCT). Reactions were run on AgPath RT-PCR (Life Technologies) kit on a QuantStudio™ 7 Flex Real-Time PCR System and the relative amounts of FAM and HEX signal used to determine the proportion of Omicron and Delta in the original samples. Validation of the assay was carried out by extracting RNA from pure Omicron and Delta supernatants, normalising by E gene copies and mixing at different ratios before measuring using the assay.

### Modelling

To quantify the efficiency with which Omicron and Delta enter hNECs by either the endosomal or TMPRSS2 pathway, we fitted a mathematical model to the data where hNECs were untreated or pretreated with Camostat, and then infected with Omicron or Delta. The mathematical model included the following processes: virus entry through either the TMPRSS2 or endosomal pathway; the eclipse phase until virus production; virus production; loss of virus infectivity; and infected cell death. The model assumed that the entry efficiency by the TMPRSS2 pathway was reduced to 0 in the presence of Camostat. Only ACE2^+^ cells could become infected by the endosomal pathway, and only ACE2^+^ TMPRSS2^+^ cells could become infected by the TMPRSS2 pathway. The proportion of ACE2^+^ and ACE2^+^ TMPRSS2^+^ cells were taken from Supplementary Table 1. The model assumed that the entry efficiency and eclipse phase duration could be different between the two pathways, and that the eclipse phase duration for the endosomal pathway was shorter or equal to that of the TMPRSS2 pathway. All model parameters were allowed to vary between virus strains. Please see Supplementary Methods for mathematical models and additional results.

**Table 1.** Virus isolates used in this study.

| Name in text | Virus name | PANGO lineage | GISAID/COG Accession | Reference |
| --- | --- | --- | --- | --- |
| WT/D614G | hCoV-19/England/IC19/2020 | B.1.13 | EPI_ISL_475572 | McKay et al<sup>49</sup> |
| Alpha | hCoV-19/England/205080610/2020 | B.1.1.7 | EPI_ISL_723001 | Brown et al<sup>50</sup> |
| Delta isolate 1 | hCoV-19/England/SHEF-10E8F3B/2021 | B.1.617.2 | EPI_ISL_1731019 | Peacock et al<sup>51</sup> |
| Delta isolate 2 | ISST987 | AY.4.2 | N/A | This study |
| Omicron/BA.1 isolate 1 | M21021166 | BA.1 | N/A | Dejnirattisai et al<sup>10</sup> |
| Omicron/BA.1 isolate 2 | hCoV-19/England/PHEP-YYNTNOA/2021 | BA.1 | EPI_ISL_7993790 | This study |
| Omicron/BA.2 | IC243335 | BA.2 | N/A | This study |

## Data availability

The sequences of the virus isolates used are available at GISAID: EPI_ISL_475572 (WT/D614G), EPI_ISL_723001 (Alpha), EPI_ISL_1731019 (Delta isolate 1), EPI_ISL_7993790 (Omicron/BA.1 isolate 2).

## Code availability

Code to reproduce all modelling results can be found at https://github.com/ada-w-yan/deltaomicron1.

## Acknowledgements and funding

The authors would like to thank Dr Matthew Turnbull and Dr Suzannah Rihn of the MRC-University of Glasgow Centre for Virus Research (CVR) for sharing their Vero E6-ACE2-TMPRSS2 cells and Gavin Screaton, Wanwisa Dejnirattisai and Alison Cowper from Oxford University for sharing the BA.1 isolate. For the other Delta and Omicron swab samples we thank Thushan de Silva at University of Sheffield, the ISARIC4C consortium and Paul Randell, Marcus Pond and colleagues at NWLP, UKHSA, and Michael Crone and Graham Taylor of Imperial College London.

This work was supported by the G2P-UK National Virology Consortium funded by the MRC (MR/W005611/1). Additional funding to DB, NT and JN were funded by The Pirbright Institute’s BBSRC institute strategic programme grant (BBS/E/I/00007038). The work at the CVR was also supported by the MRC grants (MC_UU12014/2) and the Wellcome Trust (206369/Z/17/Z). AWCY is supported by an Imperial College Research Fellowship.

## Author Contributions

T.P.P, J.C.B, J.Z., D.B. and W.S.B conceived and planned experiments. T.P.P, J.C.B, J.Z., N.T., K.S., J.N., R.K., A.W.C.Y., W.F., G.D.L., V.M.C., D.R., M.M., J.L.Q., and O.K.P. performed the experiments. T.P.P, J.C.B, J.Z., N.T., K.S., J.N., A.W.C.Y. and M.K. analysed the data. A.H.P., M.P., D.B. and W.S.B. provided supervision. T.P.P., J.C.B. and W.S.B. wrote the manuscript with input from all other authors.

## Materials and Correspondence

Correspondence to Wendy S. Barclay.

## Supplementary Figures

**Extended data Figure 1.**
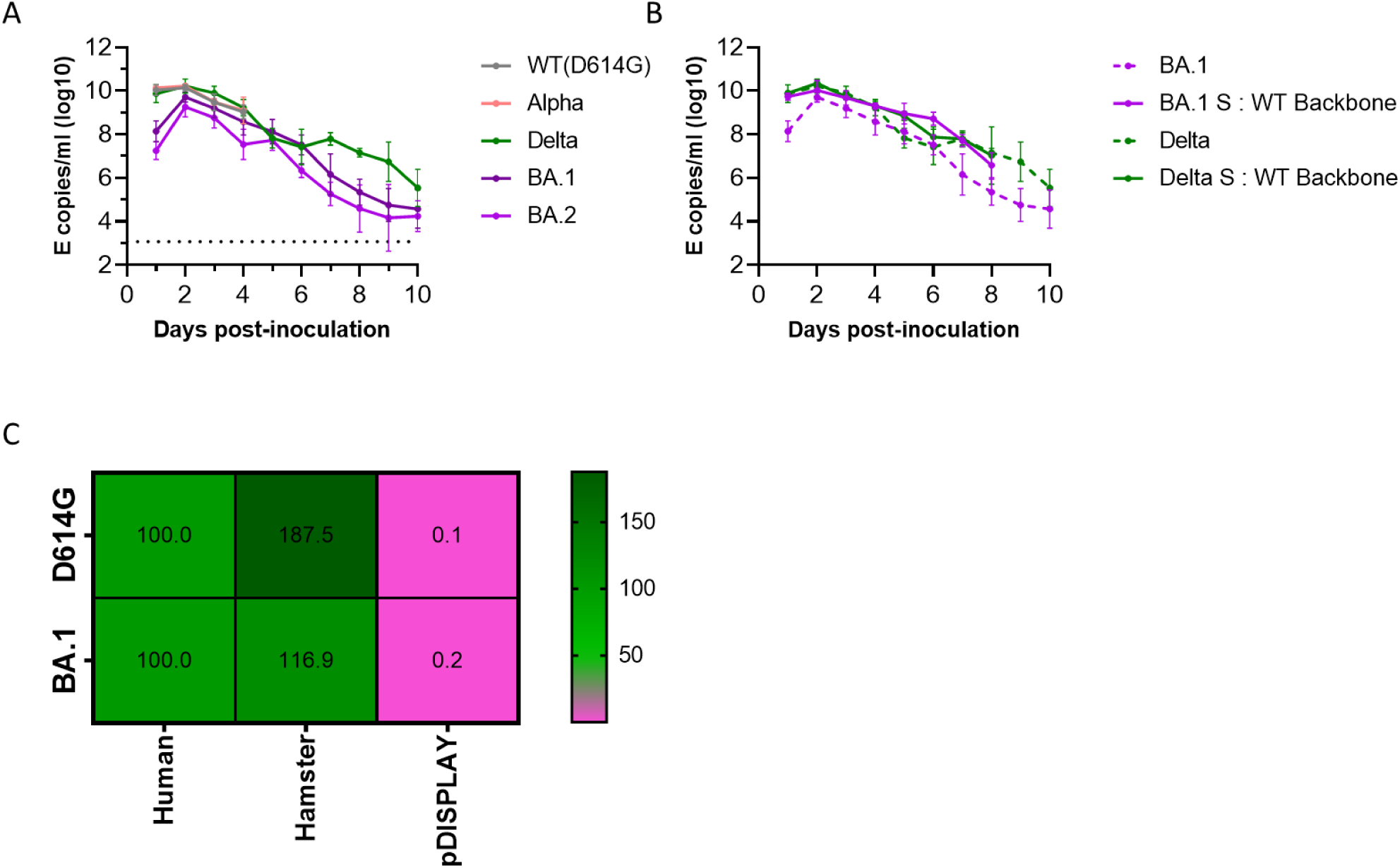
Extended data of Figure 1A,C – SARS-CoV-2 qPCR shedding of hamsters and hamster ACE2 usage. (A, B) Shedding of SARS-CoV-2 viral RNA was determined by E gene RT-qPCR on days 1-10 post-infection. (C) Receptor usage was screened using pseudoviruses expressing the indicated spike proteins into BHK-21 cells expressing the indicated ACE2 protein. Viral entry was measured by assaying luciferase activity (RLU) using the BrightGlo reagent (Promega). All assays were performed 3 independent times with mean of N=3 repeats shown.

**Extended data Figure 2.**
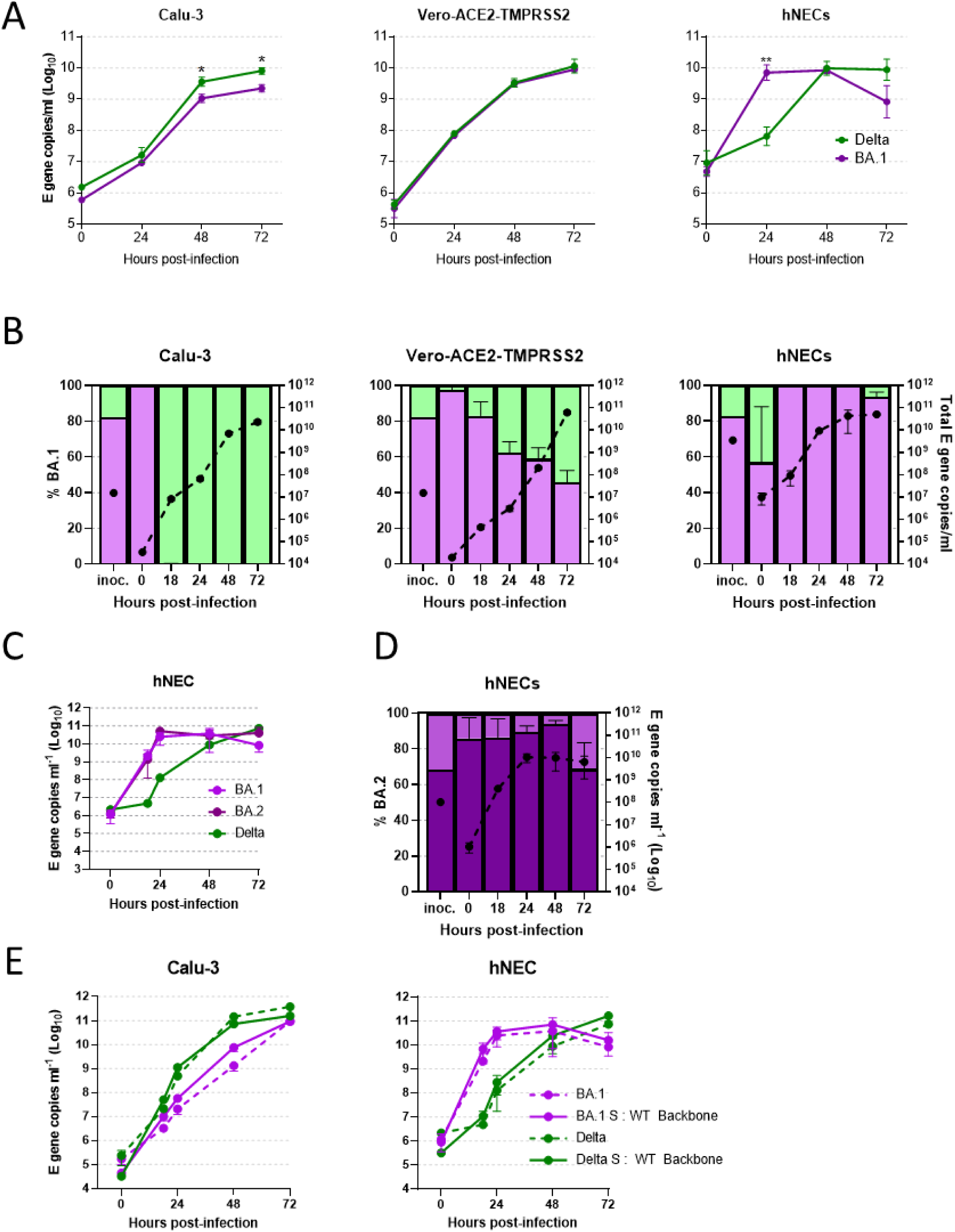
Extended data of Figure 2 – Comparative replication kinetics of Omicron and Delta. (A, C) Comparative replication kinetics of SARS-CoV-2 BA.1 and (A) Delta or (C) BA.2 variants *in vitro* and *ex vivo*. Primary human nasal epithelial cultures (hNECs) were inoculated with Delta or Omicron variants at an MOI of 0.1, and Vero-ACE2-TMPRSS2 (Vero-AT) and Calu-3 cells were inoculated at an MOI of 0.001. Infections were performed in triplicate, back titration confirmed equal inputs of the variants by E gene copies measured by RT-qPCR and infectious units measured by E gene RT-qPCR. Bars show mean + s.d.. Statistical differences measured by One-Way ANOVA on log-transformed data. *, P<0.05; **, P<0.01; ***, P<0.001; ****, P<0.0001. **–** (B,D) Competition assay between a second, independent set of SARS-CoV-2 isolates of Omicron/BA.1 (Purple) and Delta/AY.4.2 (Green) *in vitro* and *ex vivo*. A probe-based RT-qPCR assay was developed to measure the proportion of Omicron and Delta present in mixed samples (see Extended data Figure 3). The proportion of variant-specific signal is shown with the SD of 3 replicates. Equal mixtures were inoculated onto hNECs at a final multiplicity of 0.1 pfu/cell and onto Vero-AT and Calu-3 cells at an MOI of 0.001. Daily harvests were measured for total viral load by E gene qRT-PCR (black dots) and by the variant specific RT-qPCR to give variant proportions. Mean + s.d. of 3 replicates shown. (E) Comparative replication kinetics of SARS-CoV-2 Omicron and Delta isolates and chimeric spike viruses *in vitro* and *ex vivo* as shown by E gene RT-qPCR.

**Extended data Figure 3.**
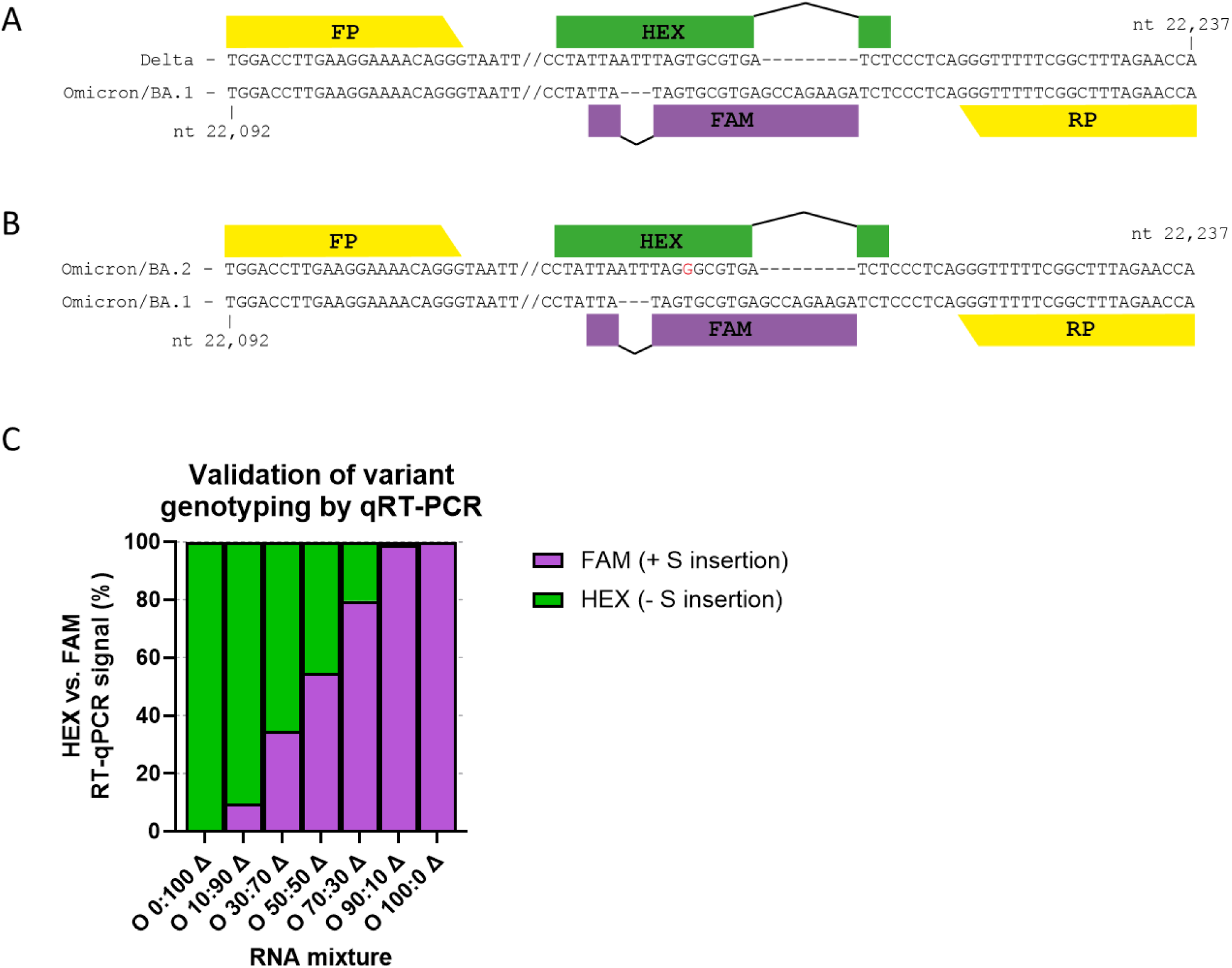
Extended data of Figure 2B - Design and validation of variant specific multiplexed RT-qPCR and independent isolate repeats. (A) Design schematic of variant specific qPCR primer probe sets for investigating the ratio of Delta to Omicron/BA.1 in competition assays. (B) Design schematic of adapted variant specific qPCR primer probe sets for investigating the ratio of BA.2 to BA.1 in competition assays. (C) Different ratios of Delta and Omicron/BA.1 RNA were mixed and the variant specific RT-qPCR probes, which specifically recognise a region of spike that includes a deletion and an insertion difference between Delta and Omicron/BA.1 were tested.

**Extended data Figure 4.**
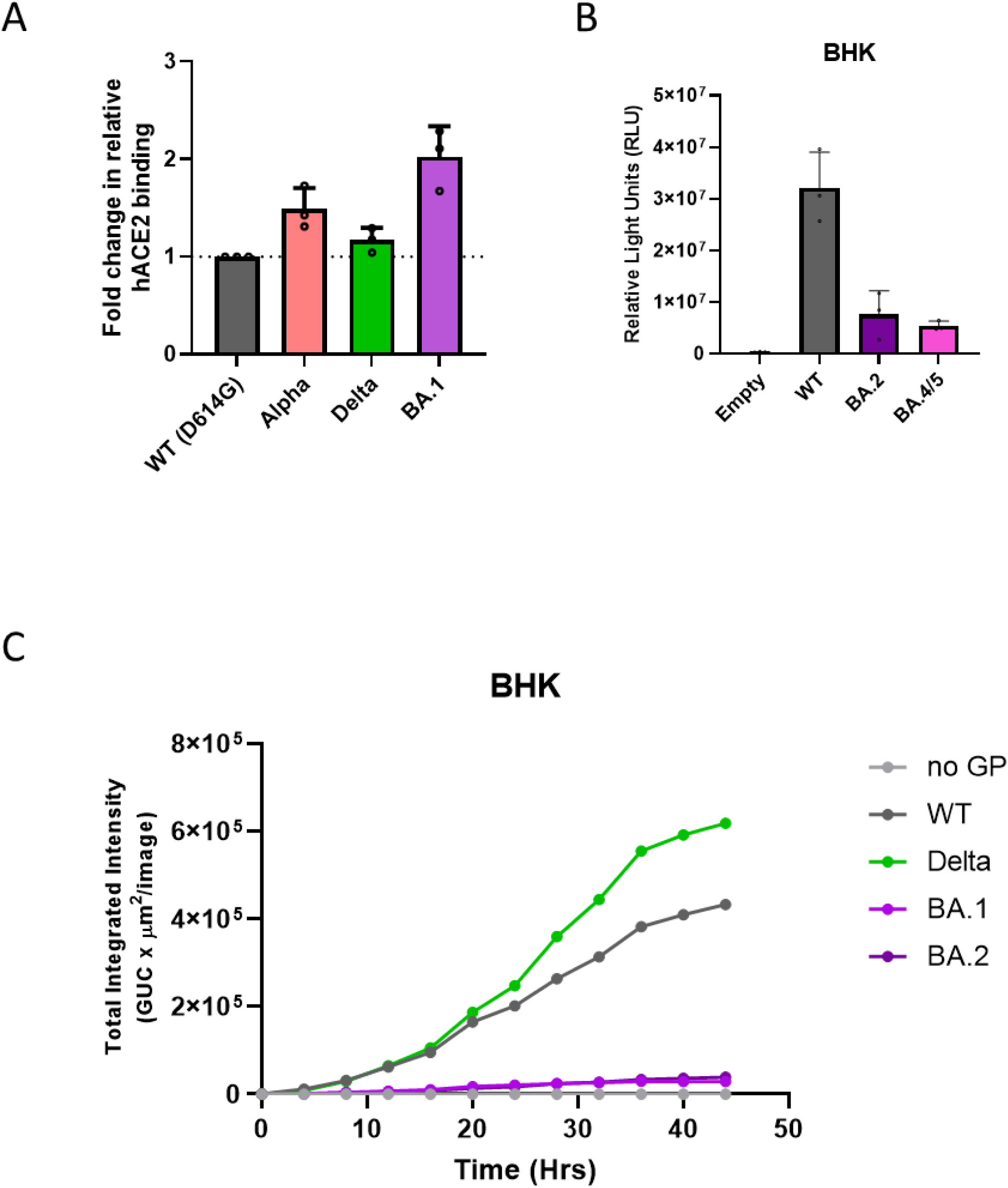
Use of common coronavirus protein receptors and receptor species specificity for SARS-CoV-2 variant entry. (A) HEK 293T cells expressing whole SARS-CoV-2 spike were incubated with recombinant hACE2-Fc(IgG) for 30 minutes, washed and incubated with Goat Anti-Human IgG Fc (DyLight^®^ 650) secondary for 30 minutes, before washing and analysis by flow cytometry. Mean + s.d. of three (N=3) independent repeats plotted. (B, C) Cell-cell fusion assays of BHK cells with rLUC-GFP1-7 expressing the stated spike protein and BHK-21 cells expressing human ACE2 and rLUC-GFP 8-11. All assays were performed in triplicate and are plotted as mean + s.d. Representative repeat of 3 independent repeats shown.

**Extended data Figure 5.**
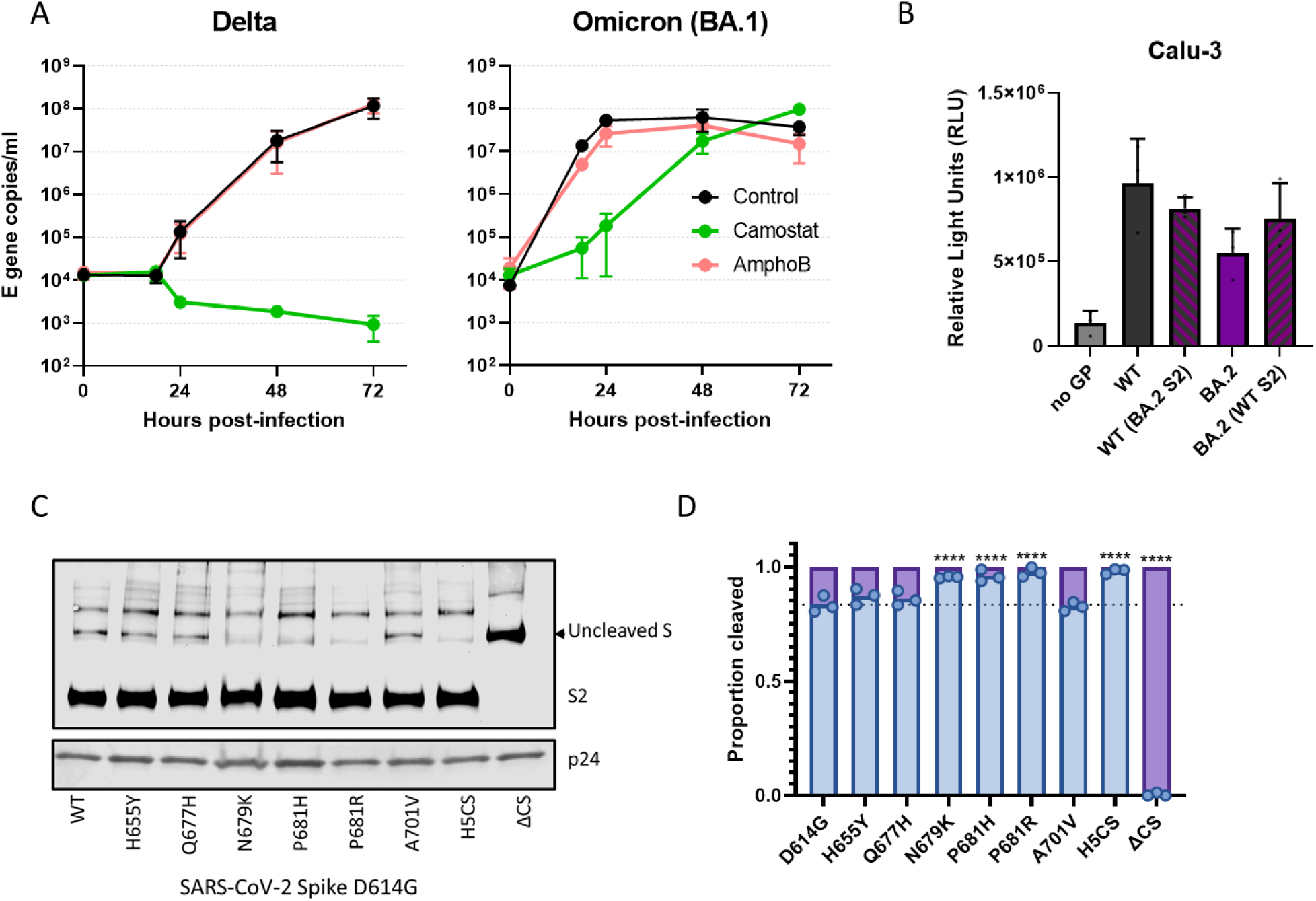
Extended data of Figure 3C. Cell entry mechanism of Omicron. Comparative replication kinetics of SARS-CoV-2 Omicron/BA.1 and Delta variants in primary human nasal epithelial cultures (hNECs) inoculated with Delta or Omicron variants at MOI of 0.1. Cells were pretreated both basolaterally and apically with either 50 µM of Camostat, 1 µM of Amphotericin B or no drug for 2 h prior to infection. 50 µM of Camostat or 1 µM Amphotericin B remained in the basolateral media for the course of the experiment. Infections were performed in triplicate and E gene copy number was determined by RT-qPCR. Plotted as mean of N=3 repeats ± s.d. Experiment performed twice with representative graph shown. (B) Cell-cell fusion assays of Calu-3 cells with rLUC-GFP1-7 expressing the stated spike protein and BHK-21 cells expressing human ACE2 and rLUC-GFP 8-11. All assays were performed in triplicate and are plotted as mean + s.d. Representative repeat shown. (C) Western blot analysis of concentrated pseudovirus expressing different SARS-CoV-2 spike mutants. Levels of lentiviral p24 shown as loading control. Representative blot of N=3 repeats. (D) Western blot densitometry of cleavage of mutants from part (C), N=3, data plotted as mean + s.d. Statistical differences measured by one-way ANOVA with multiple comparisons to WT. *, P<0.05; **, P<0.01; ***, P<0.001; ****, P<0.0001

**Extended data Figure 6.**
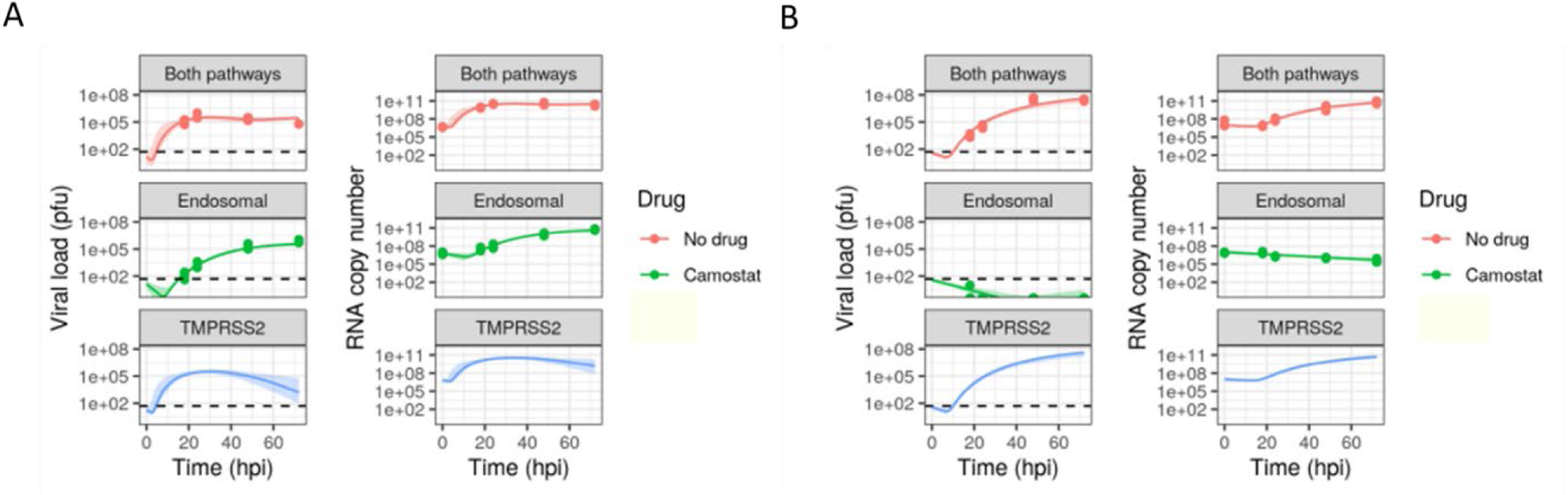
Model fits to the growth kinetics data for (A,B) Omicron or (C,D) Delta in hNECs. Dots show (A, C) the infectious viral load in plaque forming units (pfu) or (B, D) the total viral load in E gene copy numbers with no drugs (red), with Amphotericin B (yellow), or with Camostat (green). Lines show maximum likelihood model fits to the data; the blue and purple lines are the maximum likelihood model predictions for the endosomal pathway only in the absence of IFITM, and the TMPRSS2 pathway only. The shaded ribbons show 95% credible intervals for the model predictions. The dashed line shows the limit of detection.

**Supplementary Table S1.** Metaanalysis of single cell transcriptomic data sets^32-37^.

| ACE2+TMPRSS2- | ACE2-TMPRSS2+ | ACE2+TMPRSS2+ | ACE2-TMPRSS2- | ACE2+ | origin |
| --- | --- | --- | --- | --- | --- |
| 0.026687510 | 0.207526101 | 0.010887164 | 0.754899224 | 0.037574675 | nasal |
| 0.008451657 | 0.191007437 | 0.007324769 | 0.793216137 | 0.015776426 | nasal |
| 0.003723372 | 0.051638894 | 0.000610389 | 0.944027345 | 0.004333761 | nasal polyp |
| 0.021927332 | 0.369990307 | 0.025203062 | 0.582879299 | 0.047130394 | nasal |
| 0.005981810 | 0.048342794 | 0.000549350 | 0.945126045 | 0.006531160 | lung |
| 0.002380516 | 0.131538790 | 0.003418177 | 0.862662516 | 0.005798694 | lung |
| 0.000305194 | 0.03167918 | 0.000732467 | 0.967283159 | 0.001037661 | lung |
| 0.003784411 | 0.217054264 | 0.003174022 | 0.775987304 | 0.006958433 | lung |
| 0.004394799 | 0.281389245 | 0.007690899 | 0.706525056 | 0.012085698 | lung |
| 0.000671428 | 0.050479155 | 0.000915583 | 0.947933834 | 0.001587011 | lung |
| 0.00927791 | 0.254165904 | 0.00982726 | 0.726728926 | 0.019105170 | lung |

**Supplementary Table S2.** Doubling time of virus in hNECs (in hours), for the endosomal pathway, TMPRSS2 pathway, or both, as estimated by fitting a mathematical model to the data. The median estimate and 95% credible interval is reported for each value. The rightmost column shows the p-value for the doubling time being different between Omicron and Delta, for each pathway. Asterisks denote statistical significance for α = 0.05 and applying the Bonferroni correction for n = 3 comparisons.

| Pathway | Omicron | Delta | p-value |
| --- | --- | --- | --- |
| Both | 2.62 [1.69, 3.23] | 3.64 [3.20, 4.17] | 0.0101* |
| Endosomal | 7.36 [6.32, 8.59] | -16.14 [-4.37, -9.35] | < 0.0001* |
| TMPRSS2 | 2.51 [1.56, 2.99] | 3.64 [3.20, 4.17] | 0.0008* |

