## Supplementary Methods for "The altered entry pathway and antigenic distance of the SARS-CoV-2 Omicron variant map to separate domains of spike protein"

### Extended methods for SARS-CoV-2 modelling

Ada Yan

May 11, 2022

#### 1 Model

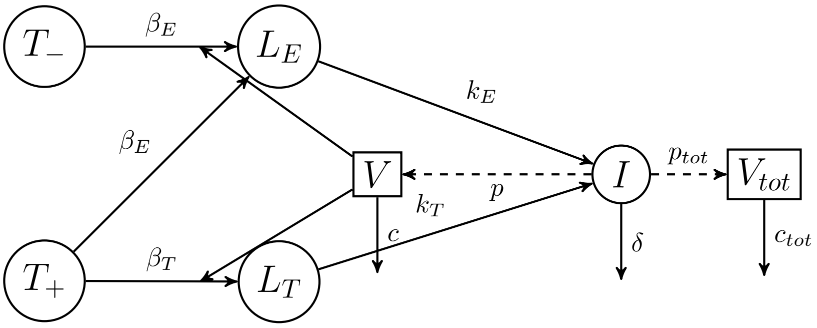


Figure 1: Compartmental diagram for model. The model also includes stages in the latent and infectious periods (not depicted).

The model equations are

$$\frac{dT_-}{dt} = -\beta_E T_- V \tag{1a}$$

$$\frac{dT_+}{dt} = -[\beta_E + \beta_T]T_+ V \tag{1b}$$

$$\frac{dL_{E1}}{dt} = \beta_E TV - n_L k_E L_{E1} \tag{1c}$$

$$\frac{dL_{Ei}}{dt} = n_L k_E (L_{E(i-1)} - L_{Ei}), i = 2, \ldots, n_L \tag{1d}$$

$$\frac{dL_{T1}}{dt} = \beta_T TV - n_L k_T L_{T1} \tag{1e}$$

$$\frac{dL_{Tj}}{dt} = n_L k_T (L_{T(j-1)} - L_{Tj}), j = 2, \ldots, n_L \tag{1f}$$

$$\frac{dI_1}{dt} = n_L k_E L_{En_L} + n_L k_T L_{Tn_L} - n_I \delta I_1 \tag{1g}$$

$$\frac{dI_m}{dt} = n_I \delta (I_{m-1} - I_m), m = 2, \ldots, n_I \tag{1h}$$

$$\frac{dV}{dt} = p \sum_{m=1}^{n_I} I_m - [c + \beta_E(1-f)(T_- + T_+) + \beta_T T_+]V \tag{1i}$$

$$\frac{dV_{tot}}{dt} = p_{tot} \sum_{m=1}^{n_I} I_m - c_{tot} V_{tot} + -[\beta_E(T_- + T_+) + \beta_T T_+]V \tag{1j}$$

The model is based on the family of target cell/infected cell/virus models, which were first applied to HIV [1], and subsequently applied to respiratory viruses [2] including SARS-CoV-2 [3]. ACE2+ TMPRSS2- target cells ($T_-$) and ACE2+ TMPRSS2+ target cells ($T_+$) are infected by virus ($V$) through the endosomal pathway at a rate $\beta_E$, while ACE2+ TMPRSS2+ target cells are also infected by virus through the TMPRSS2 pathway at a rate $\beta_T$. Cells infected by the endosomal pathway ($L_{Ei}$) and cells infected by the TMPRSS2 pathway ($L_{Tj}$) go through an eclipse phase of mean durations $k_E$ and $k_T$ respectively, before becoming cells producing virions ($I_m$). These cells produce infectious virus as measured by plaque assay ($V$) at a rate $p$, and non-infectious virus as measured by qPCR $V_{tot}$ at a rate $p_{tot}$. The mean lifetimes of cells producing virus, infectious virions, and non-infectious virus are $1/\delta$, $1/c$ and $1/c_{tot}$ respectively. The latent and infectious periods are Erlang distributed with shape parameters $n_L$ and $n_I$ respectively. This is equivalent to passing through $n_L$ and $n_I$ stages respectively, spending an exponentially distributed amount of time in each stage. Previous studies have found the latent and infectious periods of influenza virus and SHIV to be Erlang-distributed or normally distributed (which is closer to Erlang with shape parameter $> 1$ than exponential) [4, 5], hence the choice of the Erlang distribution.

The model is solved by setting $T_-(-1)$, $T_+(-1)$, $V(-1)$ and $V_{tot}(-1)$ to non-zero initial values (to be fitted), and $L_E(-1) = L_T(-1) = I(-1) = 0$. The ODEs are then solved from $t = -1$ to $t = 0$, representing a 1-hour incubation period. The inoculum is then washed off by multiplying $V(0)$ and $V_{tot}(0)$ by a constant $W$, whose value is between 0 and 1. The equations are then solved until $t = 72$ hours.

Model equations are solved using the `odin` package version 1.2.1, available at `https://github.com/mrc-ide/odin`.

The basic reproduction number for this model is

$$R_0 = \frac{[\beta_E(T_-(-1) + T_+(-1)) + \beta_T T_+(-1)]p}{\{c + [\beta_E(T_-(-1) + T_+(-1)) + \beta_T T_+(-1)]\}\,\delta}, \tag{2}$$

and the initial growth rate $r$ is computed by linearising around the disease-free equilibrium $[T_-, T_+, L_{Ei}, L_{Tj}, I_m, V] = [T_-(-1), T_+(-1), 0, \ldots, 0]$ [6, 7]. For ease of interpretation, the growth rate is converted into a doubling time using $d = \log(2)/r$.

When solving for the viral load in the presence of drugs, we assume that Camostat sets $\beta_T = 0$.

### 2 Fitting the model to data

We model the observed viral load (infectious or total) as lognormally distributed around the true viral load. For example, for a single data point for the total viral load we have

$$P(\hat{V}_{tot}(t)|\boldsymbol{\theta}) = \frac{1}{\sqrt{2\sigma_{tot}^2\pi}} \exp\left\{ -\frac{\left[\log_{10}\hat{V}_{tot}(t) - \log_{10} V_{tot}(t,\boldsymbol{\theta})\right]^2}{2\sigma_{tot}^2} \right\} \tag{3}$$

where $\hat{V}_{tot}(t)$ is the observed viral load at time $t$ and $V_{tot}(t,\boldsymbol{\theta})$ is the true viral load at time $t$ according to the model parameters $\boldsymbol{\theta}$.

In addition, for the plaque assay we model an observation threshold of $\Theta = 50$ pfu below which the viral load is treated as censored [8]. We denote below-threshold measurements as 0, so $\hat{V}(t)$ can take the value 0 or any value above (and including) $\Theta$. The likelihood of a single data point according to the plaque assay given model parameters is then given by Eq 4.

$$P(\hat{V}(t)|\boldsymbol{\theta}) = \begin{cases} \frac{1}{\sqrt{2\sigma^2\pi}} \exp\left\{-\frac{\left[\log_{10}\hat{V}(t) - \log_{10}V(t,\boldsymbol{\theta})\right]^2}{2\sigma^2}\right\} & \text{if } \hat{V}(t) \geq \Theta, \\ \int_0^{\Theta} \frac{1}{\sqrt{2\sigma^2\pi}} \exp\left\{-\frac{\left[\log_{10}x - \log_{10}V(t,\boldsymbol{\theta})\right]^2}{2\sigma^2}\right\} dx & \text{if } \hat{V}(t) = 0, \\ 0 & \text{otherwise.} \end{cases} \tag{4a}$$

Errors are assumed to be independent, so the likelihood across all data points is the product of the likelihoods of each of the data points, across the data with and without Camostat.

The compartments $V$ and $V_{tot}$ are in units pfu and RNA copy numbers, rather than concentrations pfu/mL and RNA copy number/mL which were in the raw data. This is because we assume that 1 pfu of virus and 1 target cell combine to become 1 infected cell. To convert concentrations into numbers, we multiply by the inoculum/supernatant volume 200 $\mu$ L.

The values of fixed model parameters are as follows.

| Parameter | Value | Units |
|---|---|---|
| $n_L$ | 10 | - |
| $n_I$ | 10 | - |
| $T_-(-1)$ | 8428 | cell |
| $T_+(-1)$ | 4672 | cell |
| $\frac{V_{tot}(-1)}{/}V(-1)$ | 255000 (Omicron) or 222000 (Delta) | RNA copy number / pfu |
| $\Theta$ | 50 | pfu$^{-1}$ |

Table 1: Values of fixed parameters.

The ratio $\frac{V_{tot}(-1)}{/}V(-1)$ was set to that measured in the virus stock. The initial number of target cells was calculated by multiplying the number of cells in each well, 500000, by the proportion of ACE2$^+$ TMPRSS2$^+$ or ACE2$^+$ TMPRSS2$^-$ cells in Supplementary Table 1. Only the nasal cell data in Supplementary Table 1 was used, and the proportions were averaged across the four studies.

The prior bounds for fitted parameters are as follows. Parameters with Y in the log transform column have uniform priors in log space, while the others have uniform priors in linear space. For example, the prior for $\log_{10} c$ is uniform between -2 and -0.5. For some parameters, a composite parameter rather than the original parameter is fitted, to aid identifiability. Instead of fitting $\beta_E$ and $\beta_T$, we fit $\phi_E$ and $\phi_T$. $\phi_E = \frac{\beta_E[T_-(-1)+T_+(-1)]}{\beta_E[T_-(-1)+T_+(-1)]+c}$ is the probability that one pfu successfully initiates infection in a population of susceptible cells, when only the endosomal pathway is available. $\phi_T = \frac{\beta_T T_+(-1)}{\beta_T T_+(-1)+c}$ is the probability that one pfu successfully initiates infection in a population of susceptible cells, when only the TMPRSS2 pathway is available. The endosomal pathway is known to be slower than the TMPRSS2 pathway (insert reference here), so we limit the ratio of eclipse phase rates $k_E/k_T$ to between 0 and 1. It is assumed that each infected cell produces at least one infectious virion in its lifetime, so the lower bound for the burst size $p/\delta$ is set to 1. Each infectious virion corresponds to at least one RNA copy number, so the lower bound for $p_{tot}/p$ is set to 1.

Some prior distributions were centred around values in the literature (those with citations). Others are proportions which can by definition only take values between 0 and 1 ($\phi_E$, $\phi_T$, $f$, $W$. The ratio $k_E/k_T$ is constricted to between 0 and 1 because it is assumed that the TMPRSS2 pathway is more efficient than the endosomal pathway. The range for $\delta$ was chosen based on observations of cytopathic effects 2-3 days postinfection for some of the multi-cycle growth kinetics experiments in this study. The lower bound of the ratio $p_{tot}/p$ is 1 because each plaque forming unit corresponds to at least 1 RNA copy number. The upper bound is of the order of magnitude of the ratio of RNA copy numbers to pfu in the virus stock, because we assume that $p_{tot}/p$ is similar to that for the

| Parameter | log transform | Prior bounds | Units |
|---|---|---|---|
| $c$ | Y | $[10^{-2}, 10^{-.5}]$ [?, ?] | $h^{-1}$ |
| $\phi_E$ | N | $[0, 1]$ | - |
| $\phi_T$ | N | $[0, 1]$ | - |
| $k_E$ | N | $[0, 1/72]$ [?, ?] | $h^{-1}$ |
| $k_E/k_T$ | N | $[0, 1]$ | - |
| $\delta$ | Y | $[10^{-2}, 10^{0}]$ | $h^{-1}$ |
| $p/\delta$ | Y | $[10^{0}, 10^{6}]$ [?] | pfu |
| $p_{tot}/p$ | Y | $[10^{0}, 10^{6}]$ | RNA copy number / pfu |
| $V(-1)$ | Y | | pfu |
| $W$ | N | $[0, 1]$ | - |
| $\sigma$ | N | $[0, 1]$ | - |
| $\sigma_{tot}$ | N | $[0, 1]$ | - |

Table 2: Prior bounds for fitted parameters. Parameters with Y in the log transform column have uniform priors in log space, while the others have uniform priors in linear space. For example, the prior for $\log_{10} c$ is uniform between -2 and -0.5.

process in which the virus stock was grown

The joint posterior distribution of model parameters is obtained using an adaptive Metropolis-Hastings algorithm, which we implemented in R version 4.1.2 as the package `lazymcmc`, available at `https://github.com/ada-w-yan/lazymcmc/`. Parallel tempering (as developed by Geyer *et al.* [9] and reviewed by Earl *et al.* [10]) was implemented to improve exploration of parameter space. Five parallel chains with different temperatures were used; the temperatures were calibrated as previously described in Yan *et al.* [8]. A univariate proposal distribution was used. Model fitting was performed on a high performance cluster running R version 4.1.0, while all further analysis and plotting was conducted using R version 4.1.2. To assess convergence, three such sets of five parallel chains were run, after which convergence was assessed using the `coda` package version 0.19-1 [11], as previously described in Yan *et al.* [8].

### 3 Post-fitting analysis

P-values were calculated for the doubling time for each pathway. For each pathway and cell type, the $p$-value for the doubling time being different between Omicron and Delta was computed by sampling the doubling time with replacement from each marginal posterior distribution, and letting $q$ be the proportion of sampled pairs whose ratio exceeds 1. The $p$-value was then $p = 2(min(q, 1 - q))$.

Model predictions for the infectious and total viral loads (in the absence of noise) were calculated for the following scenarios:

- Both pathways (full model);
- Endosomal pathway only ($\beta_T = 0$); and
- TMPRSS2 pathway only ($\beta_E = 0$).

The infectious and total viral loads in each scenario are sampled over the posterior distribution, and 95% credible intervals are constructed by taking the 2.5% and 97.5% percentile. The maximum likelihood trajectories are calculated separately using the maximum likelihood parameters.

Code to reproduce all results can be found at `https://github.com/ada-w-yan/deltaomicron1`.

[1] Perelson AS, Neumann AU, Markowitz M, Leonard JM, Ho DD. HIV-1 Dynamics in Vivo: Virion Clearance Rate, Infected Cell Life-Span, and Viral Generation Time. Science. 1996;271(5255):1582–1586. doi:10.1126/science.271.5255.1582.

[2] Baccam P, Beauchemin C, Macken CA, Hayden FG, Perelson AS. Kinetics of Influenza A Virus Infection in Humans. J Virol. 2006;80(15):7590–7599. doi:10.1128/JVI.01623-05.

[3] Gonçalves A, Bertrand J, Ke R, Comets E, de Lamballerie X, Malvy D, et al. Timing of Antiviral Treatment Initiation is Critical to Reduce SARS-CoV-2 Viral Load. CPT: Pharmacometrics & Systems Pharmacology. 2020;9(9):509–514. doi:https://doi.org/10.1002/psp4.12543.

[4] Holder BP, Beauchemin CAA. Exploring the effect of biological delays in kinetic models of influenza within a host or cell culture. BMC Public Health. 2011;11(Suppl 1):S10.
